# Genetic and chemical inhibition of autophagy in zebrafish induced myeloproliferation

**DOI:** 10.1101/2021.06.14.448302

**Authors:** Kazi Md Mahmudul Hasan, Xiang-Ke Chen, Zhen-Ni Yi, Jack Jark-Yin Lau, Alvin Chun-Hang Ma

**Affiliations:** Department of Health Technology and Informatics, The Hong Kong Polytechnic University, Hong Kong, China; Laboratory of Neurodegenerative Diseases, School of Biomedical Sciences, The University of Hong Kong, Pokfulam, Hong Kong, China

## Abstract

Autophagy is an evolutionary conserved and dynamic lysosomal degradation process for cellular homeostasis and remodelling, which is essential for the development and maintenance of different hematopoietic fates. However, the roles of autophagy in definitive hematopoiesis remain elusive. Here, we exploited zebrafish (*Danio rerio*) to investigate the effect of knocking-out unc-51 like autophagy activating kinase 1b and 2 (*ulk1b* and *ulk2*), homologous of human ULK1 and ULK2, respectively, on definitive hematopoiesis. Upon *ulk1b* or *ulk2* mutation, autophagosome formation was blocked in zebrafish embryos. More importantly, pan-leukocytes (*lcp1*), common myeloid progenitors (*spi1b*), neutrophils (*mpx*), and macrophages (*mpeg1*.*1*) significantly elevated, while the hematopoietic stem and progenitor cell (HSPC) (*myb*), erythroid progenitors (*gata1*), and embryonic hemoglobin (*hbae1*.*1*) significantly reduced in the caudal hematopoietic tissue (CHT) of *ulk1b* or *ulk2* mutant zebrafish embryos. On the other hand, chemically modulated autophagy induction by calpeptin, a downstream autophagy inducer for *ulk* complex, was insufficient to ameliorate dysregulated hematopoiesis in both *ulk1b* and *ulk2* mutants. Conversely, autophagy inhibitor 3-Methyladenine functioned parallelly with the *ulk* mutants to maintain defective hematopoiesis. These observations raised a link between autophagy and definitive hematopoiesis and potentiates the fact that autophagy deficiency incorporates with myeloproliferation and anemia, which warrants the significance of autophagy in regulating definitive hematopoiesis.

## Introduction

Macroautophagy/autophagy is an adaptive and highly conserved metabolic process characterized by double membrane vesicle formation (autophagosome) and lysosome-dependent degradation under various physiological and pathophysiological conditions such as cellular development and differentiation, immunity, cancer, and aging.^1-4^ Autophagy is initiated with the activation of unc-51 like autophagy activating kinase (ULK1) complex, which primarily consists of ULK1, autophagy-related protein 13 (ATG13), focal adhesion kinase family-interacting protein of 200 kDa (FIP200), ATG101, ULK2 and subsequently regulated by a series of ATGs, such as ATG5, Beclin 1 (BECN), and ATG7.^5^ In particular, both ULK1 and ULK2 have redundant roles in the canonical autophagy machinery and double knockout of ULK1/2 in mice showed neonatal mortality^6^ similar to the loss of other core autophagy genes such as ATG5,^7^ ATG7,^8^ ATG9a^9^ and ATG16L.^10^ As a kinase protein of the ULK1 complex, the ULK1 initiates the phosphorylation of Beclin-1 and vacuolar protein sorting 34 (VPS34) lipid kinase,^11^ ATG14^12^ and promotes membrane recycling via ATG9.^13, 14^ Moreover, the ULK1-FIP200 protein complex plays crucial role during the formation of starvation-induced ATG8/LC3 (microtubule-associated protein light chain 3) puncta and omegasomes on which phagophores emerge.^15^ In addition to canonical autophagy, non-canonical autophagy was also identified, which involves only a subset of ATGs.^16^ For instance, ULK1-dependent ATG5-independent non-canonical autophagy is involved in the clearance of mitochondria from reticulocytes.^17^ Besides, the LC3-associated phagocytosis (LAP), which is a non-canonical autophagy in phagocytes characterized by single-membrane vesicle formation, is ULK1-indepndent.^18^

The discovery of multiple autophagy-related genes (Atgs) in the past two decades has geared up research to understand the link between autophagy and hematopoiesis. Previous studies demonstrated that the loss of Atg7 in the mice hematopoietic systems leading to aberrant HSC functions, severe myeloproliferation,^19, 20^ anemia and lymphopenia,^21^ while as a mammalian counterpart of yeast Atg17, conditional deletion of *FIP200* in mice hematopoietic tissues resulted in HSC depletion, aberrant myeloid expansion and erythroblastic anemia.^22^ Thus, different ATGs may play distinct roles in hematopoiesis. In addition to normal hematopoiesis, autophagy also implicated in hematological malignancies. Similar to other cancers,^23, 24^ the role of autophagy in hematological malignancies is paradoxical and could be either pro-oncogenic or anti-oncogenic depending on cellular context.^25-28^ Nevertheless, the precise role of ULK1 and ULK2 dependent autophagy in the context of hematopoiesis remains unclear. In particular, the complex network and underlying mechanism of canonical and non-canonical autophagy in regulating hematopoiesis remains to be investigated.

Zebrafish (*Danio rerio*) has emerged in recent years as a robust *in vivo* model for autophagy and hematopoietic research with unique characteristics, including high fecundity, optical transparency, and feasibility for genetic and chemical manipulation. The small vertebrate has been widely used to study hematopoiesis and model human hematopoietic malignancies.^29-31^ While it remains difficult to experimentally observed autophagy in other whole-animal model organisms, zebrafish has been demonstrated as a unique vertebrate model to study autophagy *in vivo* at cellular level,^32^ highlighted the potential of using zebrafish model to study the complex role of autophagy in hematopoiesis. Here we used zebrafish *ulk1b* and *ulk2* mutant models as the paradigm of autophagy dysfunction and depicted the previously unknown role of ulk-dependent autophagy in regulating definitive myelopoiesis.

## Methods

### Zebrafish husbandry and maintenance

Wild-type, Tg(*GFP-Lc3*), and Tg(*coro1a:DsRed*) zebrafish lines were maintained under standard aquatic conditions. Fish was fed with hatched brine shrimp twice a day and embryos were collected from natural spawning and staged as described previously.^33, 34^ Protocols for microinjection and whole-mount in situ hybridization (WISH) have been described previously.^29, 35^ Conventional pigmentation inhibition with 1-phenyl 2-thiourea (PTU) was refrained to avoid unwanted autophagy induction.^32^ Fixed embryos were bleached with 1% KOH and 3% H2O2 before WISH. Live imaging was performed using Lightsheet fluorescent microscope. All animal experiments were conducted in accordance with protocols approved by the Animal Subjects Ethics Sub-Committee (ASESC) of The Hong Kong Polytechnic University.

### Generation of the *ulk1a, ulk1b* and *ulk2* mutants by TALEN

Transcription activator-like effector nucleases (TALEN) pairs targeting genomic sequence of zebrafish *ulk1a Exon-2, ulk1b Exon-4* and *ulk2 Exon-7* were designed and synthesized as described previosuly.^36, 37^ mRNA encoding *ulk1a, ulk1b* and *ulk2* TALEN pairs were *in vitro* transcribed with T3 mMessage mMachine transcription Kit (Ambion, #AM1348) and microinjected into the yolk of one-cell-stage zebrafish embryos. Mutagenic activities were confirmed by restriction fragment length polymorphism (RFLP) assay and Sanger sequencing as previously described.^38^ Stable *ulk1b* mutants (*ulk1b*^*-/-*^) and *ulk2* mutants (*ulk2*^*-/-*^) were obtained from heterozygous *ulk1b* mutants (*ulk1b*^*+/-*^) and *ulk2* mutants (*ulk2*^*+/-*^) incross respectively. We further incrossed the *ulk1b*^*+/-*^ mutants with the double transgenic Tg(*GFP-Lc3*;*coro1a:DsRed*) to obtain the heterozygous mutants and subsequently incross the heterozygous mutants to obtain the double transgenic homozygous *ulk1b* mutants [Tg(*ulk1b*^*-/-*^*:GFP-Lc3*;*coro1a:DsRed*)].

### Western blot analysis

Embryos were deyolked before mechanically homogenized in CelLytic™ MT Cell Lysis Reagent (Sigma Aldrich, #C3228). Protein concentrations were measured by BCA assay kits (Thermo Scientific™, #23225) and protein lysates were mixed with sodium dodecyl sulfate (SDS) loading buffer and heat denatured. Protein samples were then electrophoresed on 12% TGX™ FastCast™ Acrylamide Kit (Bio-Rad Laboratories, Inc., #1610175), transferred to polyvinylidene difluoride (PVDF) membrane (Bio-Rad Laboratories, #1620264) and blocked in 5% non-fat dry milk (Bio-Rad Laboratories, #1706404). Membranes were then hybridized with anti-*Lc3b* (Abcam, #ab48394; 1:1000) or anti-*GAPDH* (Cell Signaling Technology, #2118; 1:20000) at 4°C overnight. PVDF membranes were further washed in TBST and incubated with goat anti-rabbit secondary antibody (Abcam, #ab6721; 1:3000) for 2 hours at room temperature before signal development with Western ECL Substrate (Bio-Rad Laboratories, #1705061) and imaging under ChemiDoc XRS+ System (Bio-Rad Laboratories).

### Autophagy modulator treatment and LysoTracker probe staining

Embryos were treated with calpeptin (Selleckchem, #S7396), chloroquine (Selleckchem, #S4157) and 3-Methyladenine (3-MA) (Selleckchem, #S2767) at 50 µM, 100 µM and 10 mM, respectively. Fluorescent dye LysoTracker™ Red DND-99 (Invitrogen, #L7528) targeting lysosomes was diluted to a final concentration of 10µM. 4 dpf Tg(*GFP-Lc3*) zebrafish embryos were incubated with the diluted LysoTracker™ Red DND-99 in dark at 28.5 °C for 45 minutes.^39, 40^ Subsequently, embryos were rinsed 3 times with E3 fish water prior to imaging.

### Lightsheet and confocal microscopic imaging

Zebrafish embryos were anesthetized with tricaine (Sigma-Aldrich, # A5040) at a concentration of 0.164 mg/ml and mounted in 1.5% low gelling temperature agarose (Sigma-Aldrich, # A9045) into glass capillary and 35 mm glass-bottom confocal dish for light sheet and confocal imaging, respectively. Live images were acquired by Zeiss Lightsheet Z.1 Selective Plane Illumination Microscope (Carl Zeiss Microscopy, NY, USA) with a 20× objective lens and Leica TCS SPE Confocal Microscope (Leica Microsystems, Wetzlar, Germany) with the 10× and 40× objective lenses. Images were further processed and analyzed with ZEN (Carl Zeiss Microscopy, NY, USA), Leica LAS-X (Leica Microsystems, Wetzlar, Germany) imaging software and ImageJ (National Institutes of Health, USA), respectively.

### Phospho-Histone H3 (pH3) immunostaining

Tg(*mpx:EGFP*) embryos at 3 dpf were fixed with 4% paraformaldehyde (PFA) at room temperature for 4 hours and subsequently permeabilized with pre-chilled acetone at -20 °C for 20 minutes. After washing with phosphate-buffered Saline with 0.1% Tween 20 Detergent (PBST), embryos were incubated in block buffer (0.1% bovine serum albumin, 0.1% dimethyl sulfoxide (DMSO), 2% normal goat serum and 0.2% Triton-X100 in PBS) for 30 minutes and hybridized with polyclonal rabbit anti-phospho-Histone H3 (Ser10) antibody (Cell Signaling Technology, #9701; 1:1000) at overnight at 4°C. After washing with PBST, embryos were incubated with Alexa Fluor 594 goat anti-rabbit secondary antibody (Invitrogen, #A-11012) at 1:500 for an hour at room temperature before final PBST washing prior to imaging.

### Quantitative analysis and statistics

To measure the relative number of autophagosome (*GFP-Lc3*^*+*^), lysosome (LysoTracker Red^+^) and autolysosome (*GFP-Lc3*^+^ and LysoTracker Red^+^) puncta, maximum intensity projections (MIPs) were performed using Z-Stack images at the midbrain sections to monitor and quantify the relative autophagosome, lysosome and autolysosome numbers inside the neurons (20 out of 100 layers were taken to clearly monitor puncta per neuron). EGFP (green color) positive autophagosome puncta, mCherry (magenta color) positive lysosome puncta and white color positive autolysosome puncta were defined by fluorescent intensity occupying more than one pixel and distinguishable from the background signals.^41^ Overall puncta inside the neuron cells were counted from the whole midbrain section for each sample. The number of autophagosome, lysosome and autolysosome puncta per neuron cell was calculated by counting the total number of color specific puncta divided by the number of neuron cells in each embryo and at least 10 embryos were recruited to complete the triplicate. ImageJ software version 1.8.0 (NIH) was used to quantify western-blot results. Statistical analysis was performed by two-way analysis of variance (ANOVA) with Tukey’s multiple comparisons tests in the mutant and drug treated groups comparing each cell mean with every other cell mean using GraphPad Prism, version 7 (GraphPad Software, CA, USA). Mann-Whitney nonparametric U-test were applied in other analysis. In all experiments, results were presented as mean ± standard error of the mean (SEM) and P-value less than 0.05 (P < 0.05) were considered statistically significant.

## Results

### TALEN-mediated mutagenesis of zebrafish *ulk1b* and *ulk2*

In zebrafish, *ulk1* is duplicated into *ulk1a* and *ulk1b*. Phylogenetic analysis revealed that zebrafish *ulk1b* and *ulk2* are orthologues of human *ULK1* and *ULK2*^42^ (Figure S1A) with highly conserved syntenic region (Figure S1B). Zebrafish *ulk1b* and *ulk2* expressed ubiquitously during early embryonic development and later predominantly in the head region (Figure S1C and S1D). While *ulk1b* also expressed in somites at 24 hour-post-fertilization (hpf) (Figure S1C), both *ulk1b* and *ulk2* expressed in trunk neural crest cells and caudal hematopoietic tissue (CHT) at 48 hpf (Figure S1C and S1D). To investigate the role of autophagy in hematopoiesis, we targeted zebrafish *ulk1b* and *ulk2* with TALEN (Figure S2A and S2B) and somatic targeting of *ulk1b Exon-4* and *ulk2 Exon-7* were confirmed by RFLP assay (Figure S2C). Stable heterozygous F1 carried a 5-base pair (bp) and 7-bp frame-shifting deletion in *ulk1b* and *ulk2*, respectively, were identified and confirmed with Sanger sequencing, which will result in pre-mature stop and truncation of ulk1b and ulk2 (Figure S2D and S2E).

### Autophagy activation, but not autophagy flux was suppressed in *ulk1b* and *ulk2* mutants

We first examined autophagy in *ulk1b* mutant. Stable homozygous *ulk1b* mutants (*ulk1b*^*-/-*^) embryos with Tg(*GFP-Lc3*) background were stained with Lysotracker Red and the relative number of autophagosomes, lysosomes and autolysosomes were significantly decreased in *ulk1b*^*-/-*^ mutants (Figure 1A), indicated a decrease in autophagy activation. Similar results were observed in somatic *ulk2* mutant (*ulk2*^*Mut*^) though lysosome numbers were not notably affected. (Figure S3A). Western blot also confirmed the decrease in Lc3-II/GAPDH (Figure 1B and S3B).

**Figure 1.**
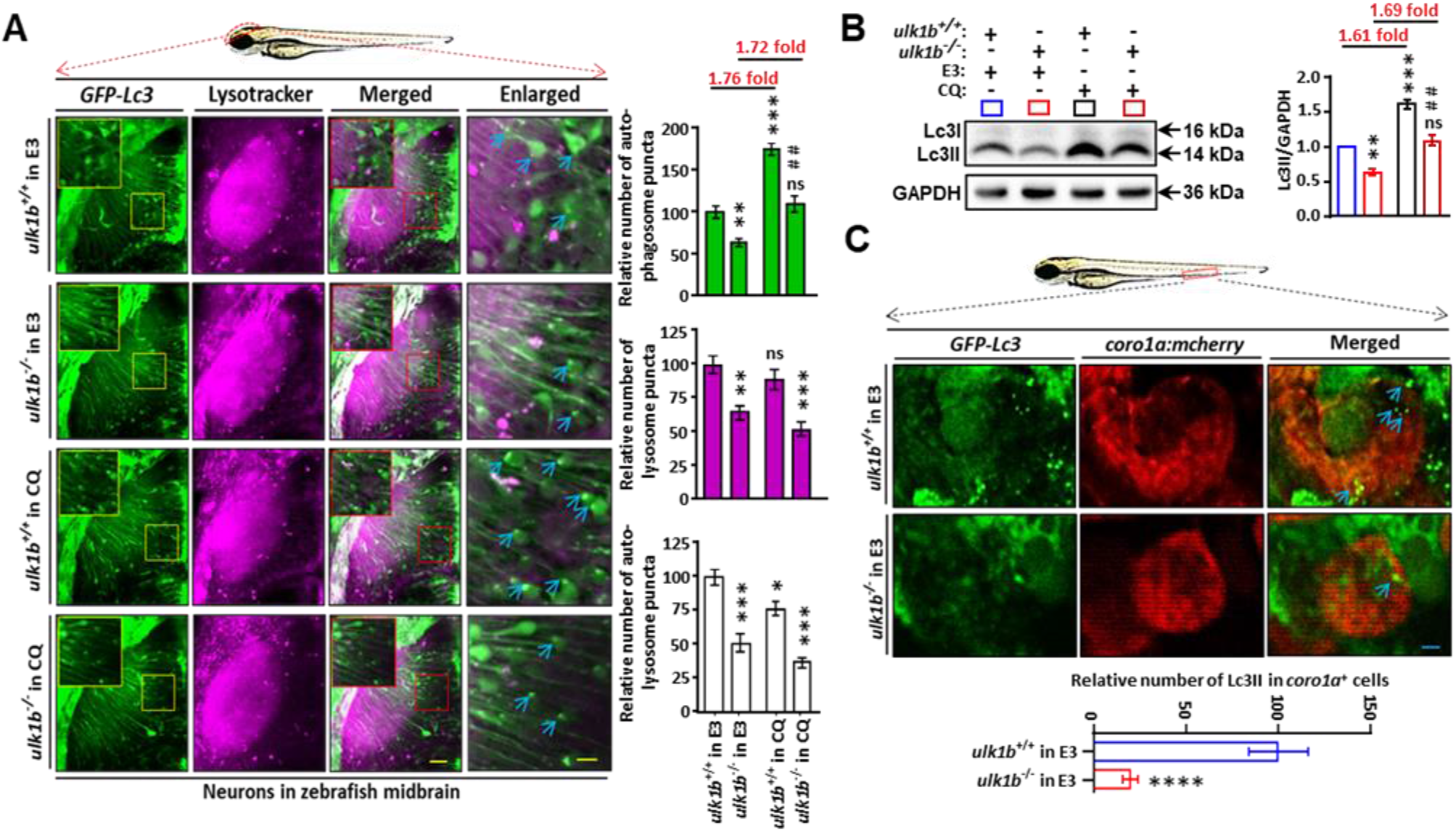
Loss of *ulk1b* inhibits autophagy activation but not autophagic flux in zebrafish. (A) Imaging of Lc3^+^ cells in the midbrain. The relative number of autophagosome (GFP-*Lc3*), lysosome (LysoTracker-red^+^) and autolysosome (merged-white^+^, fusion of GFP-*Lc3* and LysoTracker-red^+^) puncta per cell in the midbrain neuron were counted based on Z-Stack image analysis (20 layers out of 100 layers) with maximal intensity projection (MIP). All four groups (*ulk1b* ^+/+^ in E3, *ulk1b* ^-/-^ in E3, *ulk1b* ^+/+^ in CQ and *ulk1b* ^-/-^ in CQ) individually comprises in a total of at least ten Tg(GFP-*Lc3*) experimental embryos to complete three biological replicates. Red and yellow boxes indicating the autophagosome (GFP-*Lc3*) and autolysosome (merged-white^+^, GFP-*Lc3* and LysoTracker-red+) puncta respectively. Representative bar diagrams of showing the number of co-localized puncta in the neuron cell of 4 dpf drug treated and untreated zebrafish embryos. Scale bar: 40µm (Merged) and 4µm (Enlarged). (B) Western blot results showing the Lc3-II protein level of wild type siblings (*ulk1b*^*+/+*^) and homozygous mutants (*ulk1b*^*-/-*^) treated with 100µM CQ. Relative Lc3-II protein level was normalized by GAPDH while set up the *ulk1b*^+/+^ value as 1.0. Each group comprising in a total of 75 embryos for three independent experiments. Blue color box: *ulk1b*^+/+^ in E3; red color box: *ulk1b*^-/-^ in E3; black color box: *ulk1b*^+/+^ in CQ; brick red color box: *ulk1b*^-/-^ in CQ. (C) Representative images showing the myeloid cell specific autophagy in between 3 dpf double transgenic [Tg(GFP-Lc3;coro1a:mCherry)] homozygous siblings and mutant zebrafish embryos treated with E3 fish water. Scale bar: 5µm. In Panel (A) and (B), statistical analysis were performed by two-way analysis of variance (ANOVA) using Tukey’s post-hoc method and error bars were presented here as mean ± standard error of mean (SEM). \**P*<0.05, \*\**P*<0.01, \*\*\**P*<0.001, \*\*\*\**P*<0.0001, # #*P*<0.01 compared to the *ulk1b*^*-/-*^ mutants and ns: non-significant.

Concomitant treatments with late-stage autophagy inhibitor, chloroquine (CQ) was performed to examine autophagy flux. Albeit the relative number of autophagosomes were significantly decreased in both *ulk1b*^*-/t*^ and *ulk2*^*Mut*^, the fold-increase in autophagosome numbers after CQ treatment was comparable between *ulk* mutants and control, indicated that the autophagy flux was not perturbed in *ulk1b*^*-/-*^ and *ulk2*^*Mut*^ mutants (Figure 1A and S3A). Autophagy in leukocytes were also examined using Tg(*GFP-Lc3*;*coro1a:DsRed*) double transgenic line. Similar to non-hematopoietic tissues, the relative number of *lc3*-postive puncta (autophagosomes) in *coro1a*-positive cells also decreased in *ulk1b*^*-/-*^ and *ulk2*^*Mut*^ (Figure 1C and S3C).

### *ulk1b* and *ulk*2 deficiency induced myeloproliferation

The effects of *ulk1b* and *ulk*2 deficiency on definitive hematopoiesis were examined by whole-mount *in situ* hybridization (WISH). Increased expression of makers associated with myeloid lineages, including myeloid progenitor (*spi1b*), pan-leukocyte (*lcp1*) and neutrophil (*mpx*) were observed in the CHT of both *ulk1b*^*-/-*^ and stable homozygous *ulk2* (*ulk2*^*-/-*^) mutants, while the expression of markers associated with hematopoietic stem and progenitors (HSPCs) (*myb*) and erythroids (*hbae1*.*1*) were significantly decreased (Figure 2A and 2B), suggested that *ulk1b* and *ulk2* knocking-out perturbed definitive hematopoiesis, in particular, induced myeloproliferation. Similar myeloproliferation was also observed in somatic *ulk1b* (*ulk1b*^*Mut*^) and *ulk2* (*ulk2*^*Mut*^) mutants (Figure S4A and S4B). Proliferation in neutrophils was examined by immunostaining of phosphohistone H3 (pH3) in Tg(*mpx:EGFP*) at 72 hpf. In both *ulk1b*^*Mut*^ and *ulk2*^*Mut*^, the percentage of pH3-postive neutrophils significantly increased (Figure S5).

**Figure 2.**
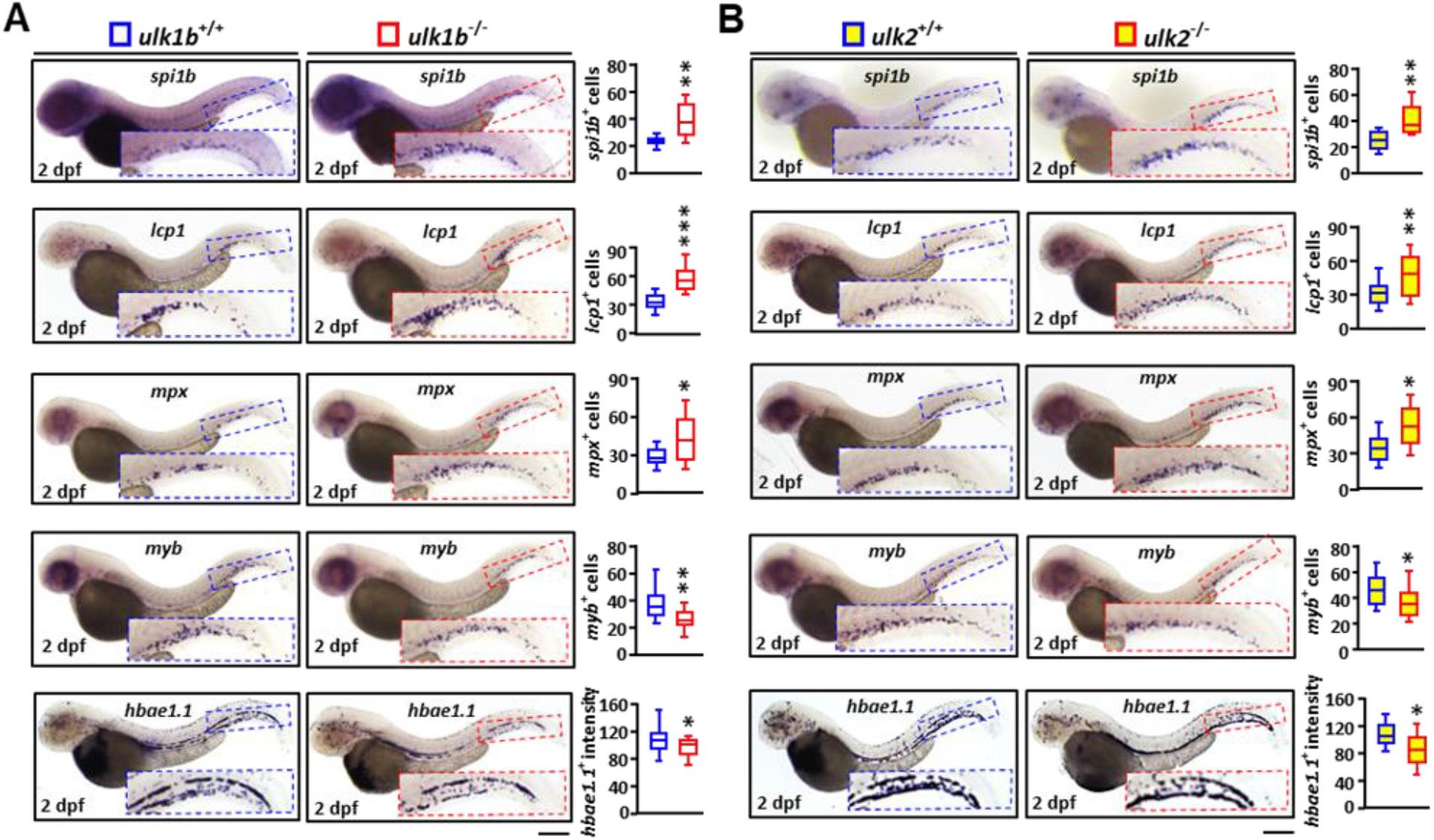
Autophagy deficient zebrafish larvae showed aberrant hematopoietic phenotypes during myelopoiesis. (A) In definitive hematapoiesis, *ulk1b* deficient homozygous mutants presented upregulation in *spi1b, lcp1* and *mpx* at 2 dpf in the CHT region compared to the wild type siblings (*ulk1b* ^+/+^) whereas *myb* and *hbae1*.*1* expressions were remarkably decreased at similar time points. (B) Similarly, in the *ulk2* mutants, *spi1b, lcp1* and *mpx* expressions in the CHT region significantly upregulated compared with the wild type siblings whereas *myb* and *hbae1*.*1* decreased. Scale bar: 300µm. In Panel (A) and (B), statistical analysis were performed by Mann-Whitney U test. All the whiskers, boxes, and central lines were shown as minimum-to-maximum values, 25th-to-75th percentile, and the 50th percentile (median), respectively. \**P*<0.05, \*\**P*<0.01, \*\*\**P*<0.001.

### Treatment with autophagy modulators also perturbed definitive hematopoiesis

To examine if the hematopoietic phenotypes observed in *ulk1b*^*-/-*^ and *ulk2*^*-/-*^ were autophagy-dependent, treatment with autophagy modulators, including 3-MA and calpeptin (autophagy inducer) were performed. Autophagy activation was significantly suppressed in embryos treated with 3-MA as shown by the decreased number of autophagosomes and autolysosomes (Figure 3A). Similar to *ulk1b* and *ulk2* mutants, treatment with 3-MA also induced myeloproliferation as shown by the increase in *lcp1* and *mpx* (Figure 3B). Expression of *myb* were markedly decreased while *hhabe1*.*1* were unaffected (Figure 3B). In contrary, calpeptin-treatment significantly increased the number of autophagosomes and autolysosomes in zebrafish embryos (Figure 4A). Also opposite to 3-MA treatment, calpeptin significantly decreased the expression of *lcp1* and *mpx*, while the expression of *myb* and *hbae1*.*1* were significantly increased (Figure 4B and Figure S6). More importantly, both *ulk1b* and *ulk2* knock-out completely reverted the effects of calpeptin treatment on autophagy and hematopoiesis (Figure 4B and Figure S6).

**Figure 3.**
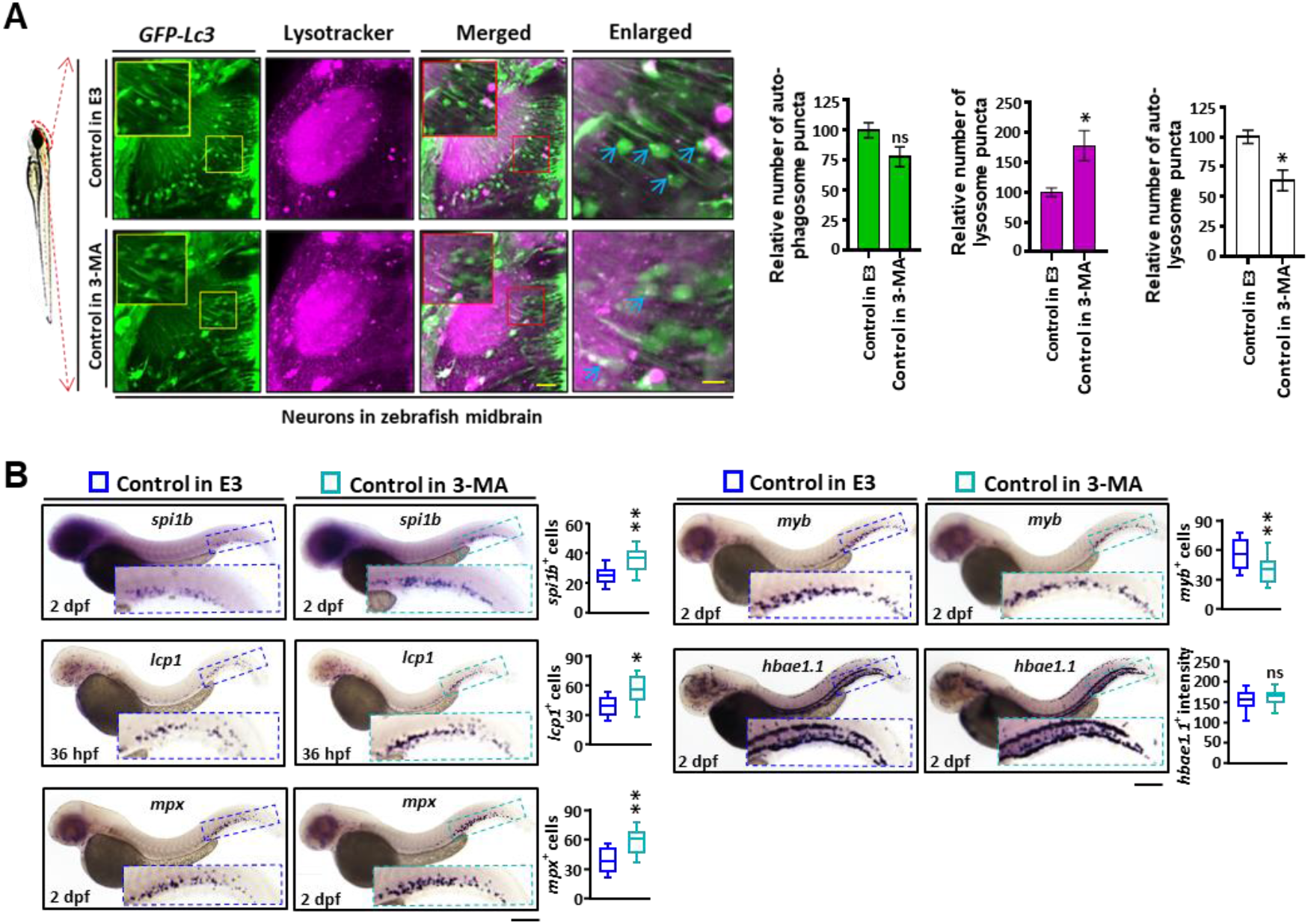
3-MA perturbed autophagosome formation and myelopoiesis. (A) Schematic illustration showing the imaging of Lc3^+^ cells in the midbrain section. The relative number of autophagosome (GFP-*Lc3*), lysosome (LysoTracker-red^+^) and autolysosome (merged-white^+^, GFP-*Lc3* and LysoTracker-red^+^) puncta per cell in the midbrain neuron were counted based on Z-stack image analysis (20 layers out of 100 layers) with maximal intensity projection (MIP). Both 3-MA treated and untreated groups individually comprises in a total of at least ten Tg(GFP-*Lc3*) experimental embryos to complete three biological replicates. Yellow and red dash boxes showing the autophagosome (GFP-*Lc3*) and autolysosome (merged-white^+^, GFP-*Lc3* and LysoTracker-red^+^) puncta respectively. 3-MA: 3-Methyladenine. Representative bar diagrams of showing the number of co-localized puncta in the neuron of 4 dpf 3-MA treated and untreated zebrafish embryos in the midbrain section. (B) In situ hybridization indicated that treatment with early autophagy inhibitor 3-MA significantly disrupted definitive hematopoiesis in the control siblings including aberrant expression of *lcp1* and *mpx* as well as ectopic downregulation of *myb* and *hbae1*.*1* in the CHT. hpf: hours post fertilization, dpf: days post fertilization, Scale bar: 300 µm. Scale bar: 40µm (Merged) and 4µm (Enlarged). In Panel (A) and (B), statistical analysis were performed by Mann-Whitney U test. In panel (A), error bars were presented here as mean ± standard error of mean (SEM) and in panel (B), all the whiskers, boxes, and central lines were shown as minimum-to-maximum values, 25^th^-to-75^th^ percentile, and the 50^th^ percentile (median), respectively. \**P*<0.05, \*\**P*<0.01 and ns, non-significant.

**Figure 4.**
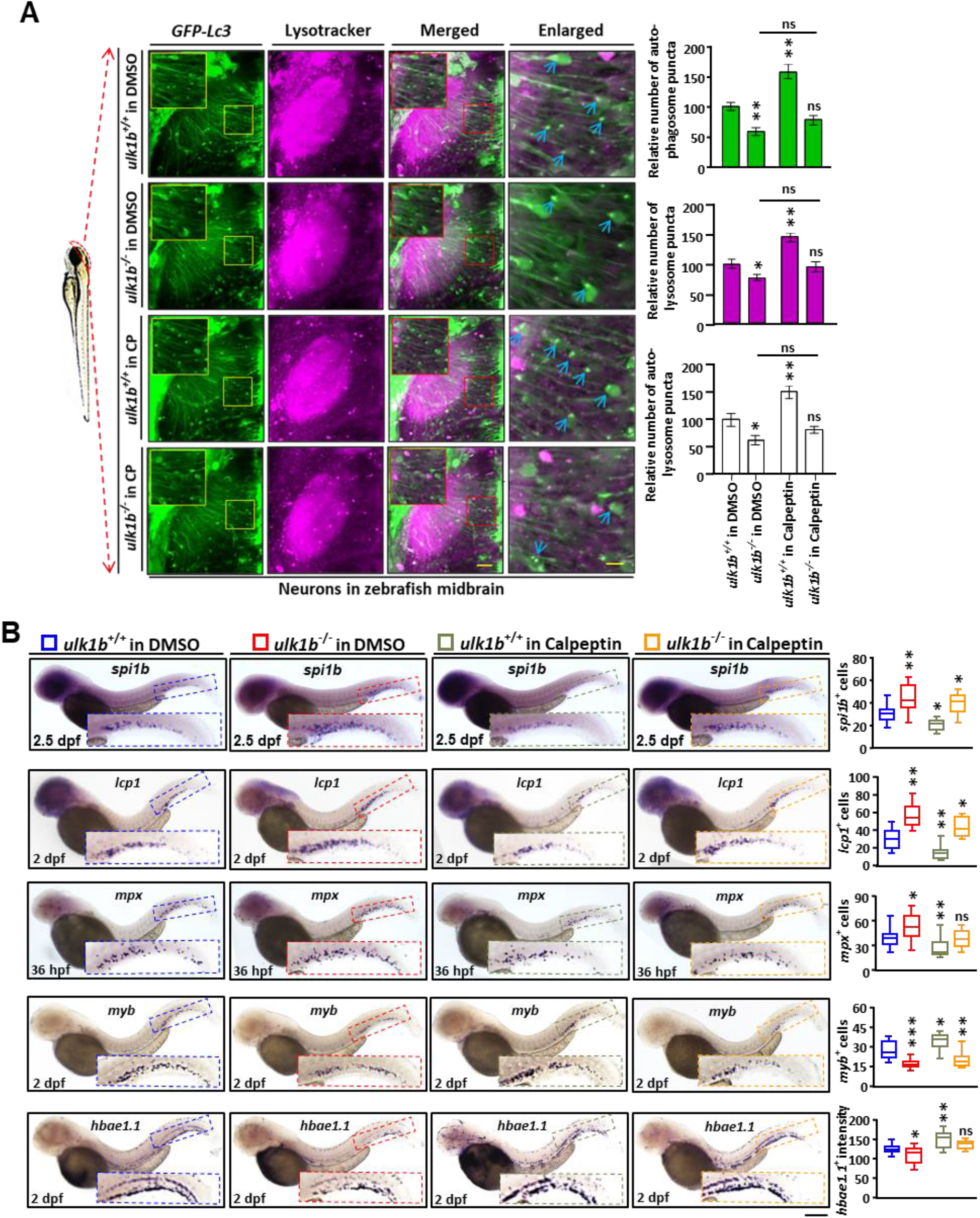
Calpeptin can partially recapitulate autophagy and ameliorate defective hematopoiesis in zebrafish mutants. (A) Schematic illustration showing the imaging of Lc3^+^ cells in the midbrain section. The relative number of autophagosome (GFP-Lc3), lysosome (LysoTracker-red^+^) and autolysosome (merged-white^+^, fusion of GFP-Lc3^+^ and LysoTracker-red^+^) puncta per cell in the midbrain neuron were counted based on Z-Stack image analysis (20 layers out of 100 layers) with maximal intensity projection (MIP). All four groups (*ulk1b*^*+/+*^ in DMSO, *ulk1b*^*-/-*^ in DMSO, *ulk1b*^*+/+*^ in calpeptin and *ulk1b*^*-/-*^ in calpeptin) individually comprises in a total of at least ten Tg(GFP-Lc3) experimental embryos to complete three biological replicates. Yellow and red color boxes indicating the autophagosome (GFP-Lc3^+^) and autolysosome (merged-white^+^) puncta respectively. Representative bar diagrams of showing the number of co-localized autophagosome, lysosome and autolysosome puncta in the neuron cell of 4 dpf calpeptin treated and untreated zebrafish embryos. Scale bar: 40µm (Merged) and 4µm (Enlarged). (B) In situ hybridization showed that calpeptin can partially rescue the *ulk1b* mutant’s *lcp1, mpx, myb* and *hbae1*.*1* phenotypes compared to the wild type siblings at 2 dpf. Scale bar: 300µm. In panel (A), statistical analysis was performed by two-way analysis of variance (ANOVA) using Tukey’s post-hoc method and error bars were presented here as mean ± standard error of mean (SEM). In panel (B), the whiskers, boxes, and central lines were shown as minimum-to-maximum values, 25^th^-to-75^th^ percentile, and the 50^th^ percentile (median), respectively. \**P*<0.05, \*\**P*<0.01 and \*\*\**P*<0.001 and ns, non-significant.

## Discussion

In this study, we took advantages of the optically clear and robust zebrafish model to investigate the role of autophagy in hematopoiesis. To model defective autophagy, we genetically targeted zebrafish *ulk1b* and *ulk2* with TALEN as described previously.^43^ Zebrafish *ulk1* is duplicated into *ulk1a* and *ulk1b* and phylogenetic analysis revealed that *ulk1b* is the orthologous of human *ULK1* and *ulk1a* knockout did not significantly affect autophagy in zebrafish embryos (data not shown). Both *ulk1b* and *ulk2* expressed ubiquitously during early embryonic development and later predominantly in the head region but also in neural crest cells, somites and CHT. The expression pattern highlighted the importance of autophagy during early embryogenesis as well as the development of neural crest cells, and explained the high autophagy level observed in skin, eye, brain and muscle cells.^32^ Both somatic *ulk1b*^*Mut*^ and *ulk2*^*Mut*^ as well as homozygous *ulk1b*^*-/-*^ and *ulk2*^*-/-*^ mutants survived with normal gross development probably due to the functional redundancy between Ulk1 and Ulk2.^44^

Defective autophagy was confirmed in both stable homozygous *ulk1b*^*-/-*^ and somatic *ulk2*^*Mut*^ mutants by the significantly decreased number of autophagosome and autolysosome. Treatment with CQ, which blocks the fusion of autophagsome and lysosome in the late stage of autophagy^45^, did not rescued the decrease in autophagosome and autolysosome numbers observed in *ulk1b*^*-/-*^ and *ulk2*^*Mut*^, suggested the blockage in autophagosome formation during early autophagy activation. While early autophagy activation was suppressed, autophagy flux was not significantly affected, which could be explained by the function of ULK1 complex in autophagy activation.^46^ Consistent with the expression of *ulk1b* and *ulk2* in CHT, autophagy puncta was also detected in *coro1a*-positive myeloid cells and more importantly, significantly decreased upon genetic targeting of *ulk1b* and *ulk2*, suggested that autophagy involves in definitive hematopoiesis, particularly in myelopoiesis.

We then examined the effects of *ulk1b* and *ulk2* mutations in definitive hematopoiesis by WISH. In particular, expansion in myeloid lineages in the expenses of HSPC and erythroid lineages were observed in homozygous mutants of *ulk1b* and *ulk2*, demonstrated the role of Ulk1 complex in regulating definitive hematopoiesis. Interestingly, aberrant proliferation in *mpx*-expressing neutrophils was also observed in *ulk* mutant models, which is similar to the phenotypes in *Atg5* knock-out mice previously reported with increased and rapid production of neutrophils^47^ and suggested Ulk and Atg5-dependent autophagy may limit immature neutrophil. To investigate if the hematopoietic phenotypes observed in *ulk1b* and *ulk2* mutants were autophagy-dependent, we chemically modulated autophagy with 3-MA and calpeptin. While treatment with 3-MA, which is a PI3K inhibitor blocking early autophagosome formation, recapitulated the myeloproliferation and treatment with autophagy inducer, calpeptin, suppressed myeloid lineages. In particular, genetic targeting of *ulk1b* and *ulk2* completely abrogated the effects of calpeptin treatment on autophagy and myeloid lineages, suggested that the ulk-deficiency induced myeloproliferation is autophagy-dependent. Nevertheless, further investigation is needed to confirm if Ulks exhibit non-canonical or autophagy-independent regulation on hematopoiesis.

During selective and nonselective autophagy, both ULK1 and ULK2 play critical roles in response to mitochondrial damage, infections and metabolic stresses in mammalian cells.^48, 49^ Likely to the ATG5 and ATG7 deficient in vivo mice models of hematopoietic systems, where all of which mediate the depletion of autophagosome or Lc3-IIB puncta in the hematopoietic tissue under basal physiologic conditions,^50, 51^ here, upon genetic and chemical targeting of autophagy, we demonstrated that *ulk1b* and *ulk2* are also important to maintain “basal” autophagy and normal hematopoiesis in zebrafish embryos. Interestingly, hematopoietic ablation of *Ulk1* in β-thalassemic mice exacerbates disease phenotypes whereas *Atg5* knockout has comparatively minor effects indicating a Ulk1 dependent and Atg5 independent role during hematopoiesis.^52^ Nevertheless, further targeting multiple and major autophagy genes either alone or in combination, downstream to the ULK1 complex, regulating different steps of the canonical pathway would further clarify the precise roles of ULK1 and ULK2 in hematopoiesis.

Autophagy has been reported to implicate in hematological malignancies. While many studies have reported that autophagy inhibition overcomes drug resistance^53, 54^ and suppresses leukemia cell growth,^55, 56^ others have demonstrated that autophagy suppression is important for leukemia development.^57, 58^ These contradictive results suggested that autophagy plays paradoxical roles in leukemogenesis depending on cellular contexts. Our results showing that autophagy suppression would lead to deregulation of normal myelopoiesis, demonstrating the important anti-oncogenic role of autophagy during early stage of myeloid malignancies. Further investigation with bona fide models of hematological malignancy and tissue specific knock-out of autophagy from the myeloid cell are warranted. For instance, zebrafish has emerged as an important model organism for human cancer and models of hematological malignancies including myeloproliferative neoplasm (MPN), acute myeloid leukemia (AML), chronic myeloid leukemia (CML) and acute lymphoblastic leukemia (ALL) with highly conserved oncogenic pathways and pharmacologic responses were reported.^59-63^ With the well-developed methodologies in autophagy study^32, 39, 64^ and genetic engineering,^65^ zebrafish could be developed into a unique modelling platform to define the role of autophagy in hematological malignancies, which will provide important information for the development of autophagy-related therapeutic strategies against these heterogeneous diseases.

## Acknowledgments

Zebrafish maintenance was supported by Fish Model Translational Research Laboratory (HTI, PolyU). Microscopic imaging was supported by University Research Facility in Life Sciences (ULS, PolyU). Transgenic zebrafish lines Tg(*GFP-Lc3*) and Tg(*coro1a:mCherry*) were kindly provided by Dr. X Xu (Mayo Clinic, Rochester, MN) and Prof. Z Wen (HKUST, HK, China), respectively. This work is supported by the FHB HMRF [03143765] and FHB HMRF [06173226] to ACM.

## Notes

### Competing Interest Statement

The authors have declared no competing interest.

