## Supplementary figures for "Genetic and chemical inhibition of autophagy in zebrafish induced myeloproliferation"

**Figure S1. Homology and developmental mRNA expression pattern of zebrafish *ulk1b* and *ulk2* genes.** (A) The genomic loci of human *ULK1* and *ULK2* surrounding q24.33 and 17p11.2 positions in chromosome 12 and 17 are syntenic with the zebrafish *ulk1b* and *ulk2* in chromosome 21 and 15 respectively. (B) Autophagy related gene *ulk1b* and *ulk2* in zebrafish (*Dr*) are evolutionary related with the human (*Hs*) autophagy related gene *ULK1* and *ULK2*. Scale bar inferring phylogenies. (C) Representative lateral views indicating the spatial expression pattern of the zebrafish *ulk1b* at one to two cell zygotes at 20 mpf (i), oblong to sphere at 4 hpf (ii) somite at 24 hpf (iii) and eye, migratory neural crest cells, head and CHT at 48 hpf (iv) stages. (D) Zebrafish *ulk2* mRNA was detected in two-cell zygote at 15 mpf (i), 50% epiboly at 6 hpf (ii), and eye, migratory neural crest cells, head and CHT at 48 hpf (iii) stages.

**Figure S2. Generation of zebrafish *ulk1b* and *ulk2* mutants by TALEN.** (A and B) Schematic illustration representing transcription activator-like effector nucleases (TALENs) mediated genome editing of zebrafish *ulk1b* gene at exon 4 and *ulk2* gene at exon 7. 15bp spacer region containing the mutation initiation site. *FokI* endonuclease cleave the target DNA fragment. *NheI* and *BamHI* restriction enzymes used in this genotyping assay to cut in a specified location of *ulk1b* and *ulk2* respectively. (C) Genotyping done by the restriction fragment length polymorphism (RFLP) assay. bp: base pair; M: marker; $ulk1b^{L\text{-}Ctrl}$: *ulk1b* left arm injected control; $ulk1b^{Mut}$: *ulk1b* TALEN mRNA injected mutant; $ulk2^{L\text{-}Ctrl}$: *ulk2* left arm injected control; $ulk2^{Mut}$: *ulk2* TALEN mRNA injected mutant. (D and E) A 5bp and 7bp deletion (Δ) mutations were confirmed by Sanger sequencing in zebrafish *ulk1b* exon 4 and *ulk2* exon 7, black dash indicating the TALEN target site to induce possible mutation. Computer aided ExPASy translator tool given both ulk1b and ulk2 truncate protein sequence corresponding to 92 amino acids (a.a) with a mutation position at Y81 residues and 62 a.a with a mutation position at S218 residues respectively.

**Figure S3. Zebrafish *ulk2* mutants inhibit autophagosome formation without affecting autophagic flux.** (A) Schematic illustration highlighted with *ulk2* showing the imaging of Lc3$^{+}$ cells in the midbrain section. The relative number of autophagosome (GFP-*Lc3*), lysosome (LysoTracker-red$^{+}$) and autolysosome (merged-white$^{+}$, fusion of GFP-*Lc3* and LysoTracker-red$^{+}$) puncta per cell in the midbrain neuron were counted based on Z-Stack image analysis (20 layers out of 100 layers) with maximal intensity projection (MIP). All four groups ($ulk2^{\text{L-Ctrl}}$ in E3, $ulk2^{\text{Mut}}$ in E3, $ulk2^{\text{L-Ctrl}}$ in CQ and $ulk2^{\text{Mut}}$ in CQ) individually comprises in a total of at least ten Tg(GFP-*Lc3*) experimental embryos to complete three biological replicates. Red and yellow boxes showing the autophagosome and autolysosome puncta respectively. L-Ctrl: TALEN left arm injected control; Mut: mutant; CQ: Chloroquine. Representative bar diagrams of showing the number of co-localized puncta in the neuron of 4 dpf CQ treated and untreated zebrafish embryos in the midbrain section. Scale bar: 40µm (Merged) and 4µm (Enlarged). (B) Western blot results showing the Lc3-II protein level wild type sibling ($ulk2^{+/+}$) and homozygous mutant ($ulk2^{-/-}$) groups treated with 100µM CQ. Relative Lc3-II protein level was normalized by GAPDH while set up the $ulk1b^{+/+}$ value as 1.0. Each group comprising in a total of 75 embryos for three independent experiments. Blue color box: $ulk2^{\text{L-Ctrl}}$ in E3; red color box: $ulk2^{\text{Mut}}$ in E3; black color box: $ulk2^{\text{L-Ctrl}}$ in CQ; brick red color box: $ulk2^{\text{Mut}}$ in CQ. (C)

Representative images showing the myeloid cell specific autophagy in between 3 dpf double transgenic [Tg(GFP-*Lc3*;*coro1a*:mCherry)] wild type siblings and *ulk2* mutant zebrafish embryos treated with E3 fish water. Scale bar: 5µm. In panel (A) and (B), statistical analysis were performed by two-way analysis of variance (ANOVA) using Tukey's post-hoc method and error bars were presented here as mean ± standard error of mean (SEM). In panel (C), data was presented as column plot. **P*<0.05, ***P*<0.01 and ****P*<0.001 and ns, non-significant.

**Figure S4. Somatic knockout of zebrafish *ulk1b* and *ulk2* perturbed definitive hematopoiesis.** (A) Whole mount in-situ hybridization (WISH) analysis showing the significant upregulation of *spi1b*, *lcp1* and *mpx* while *myb* positive cells and *hbae1.1* expression were not significantly affected in the somatic mutants ($ulk1b^{Mut}$) compared to the TALEN left arm injected control ($ulk1b^{L\text{-}Ctrl}$) at 2 dpf. Scale bar: 300µm. (B) Representative WISH images showing the significant upregulation of *spi1b*, *lcp1* and *mpx* while *myb* positive cells and *hbae1.1* expression expression were not significantly affected in the somatic mutants ($ulk2^{Mut}$) compared to the TALEN left arm injected control ($ulk2^{L\text{-}Ctrl}$) at definitive waves. Scale bar: 300µm. In both panel, statistical analysis were performed using Mann–Whitney U test. All boxes and central lines were shown as minimum-to-maximum values, $25^{th}$-to-$75^{th}$ percentile, and the $50^{th}$ percentile (median), respectively. **P*<0.05, ***P*<0.01 and ns, non-significant.

**Figure S5. Autophagy deficiency causes aberrant myeloproliferation in definitive hematopoiesis.** Double staining of neutrophil marker *mpx* (A) with phospho histone 3 (pH3) protein (B) in the CHT of 3 dpf control siblings (A-C) and $ulk1b^{Mut}$ (D-F) and $ulk2^{Mut}$ mutants (G-I). Arrowhead in figure (C), (F) and figure (I) showed *mpx* and pH3 merged puncta. Scale bar: 10µm. ***P*<0.01, *****P*<0.0001.

**Figure S6. Zebrafish $ulk2^{-/-}$ mutant's defective hematopoietic phenotypes were partially rescued by calpeptin.** (A) Schematic illustration highlighted with *ulk1b* showing the imaging of $Lc3^{+}$ cells in the midbrain section. The relative number of autophagosome (GFP-*Lc3*), lysosome (LysoTracker-$red^{+}$) and autolysosome (merged-$white^{+}$, fusion of GFP-*Lc3* and LysoTracker-$red^{+}$) puncta per cell in the midbrain neuron were counted based on Z-Stack image analysis (20 layers out of 100 layers) with maximal intensity projection (MIP). All four groups ($ulk2^{L\text{-}ctrl}$ in DMSO, $ulk2^{Mut}$ in DMSO, $ulk2^{L\text{-}ctrl}$ in calpeptin and $ulk2^{Mut}$ in calpeptin) individually comprises in a total of at least ten Tg(GFP-*Lc3*) experimental embryos to complete three biological replicates. Red and yellow boxes indicating the autophagosome (GFP-*Lc3*) and autolysosome (merged-$white^{+}$, GFP-*Lc3* and LysoTracker-$red^{+}$) puncta respectively. Representative bar diagrams of showing the number of co-localized autophagosome, lysosome and autolysosome puncta in the neuron cell of 4 dpf calpeptin treated and untreated zebrafish embryos. Scale bar: 40µm (Merged) and 4µm (Enlarged). (B) In situ hybridization showed that calpeptin can partially rescue the *ulk2* mutant's *lcp1, mpx, myb* and *hbae1.1* phenotypes compared to the wild type siblings at 2 dpf. Scale bar: 300µm. In panel (A), statistical analysis was performed by two-way analysis of variance (ANOVA) using Tukey's post-hoc method and error bars were presented here as mean ± standard error of mean (SEM). In panel (B), the whiskers, boxes, and central lines were shown as minimum-to-maximum values, $25^{th}$-to-$75^{th}$ percentile, and the $50^{th}$ percentile (median), respectively. **P*<0.05, ***P*<0.01 and ns, non-significant.

**Figure S1.**

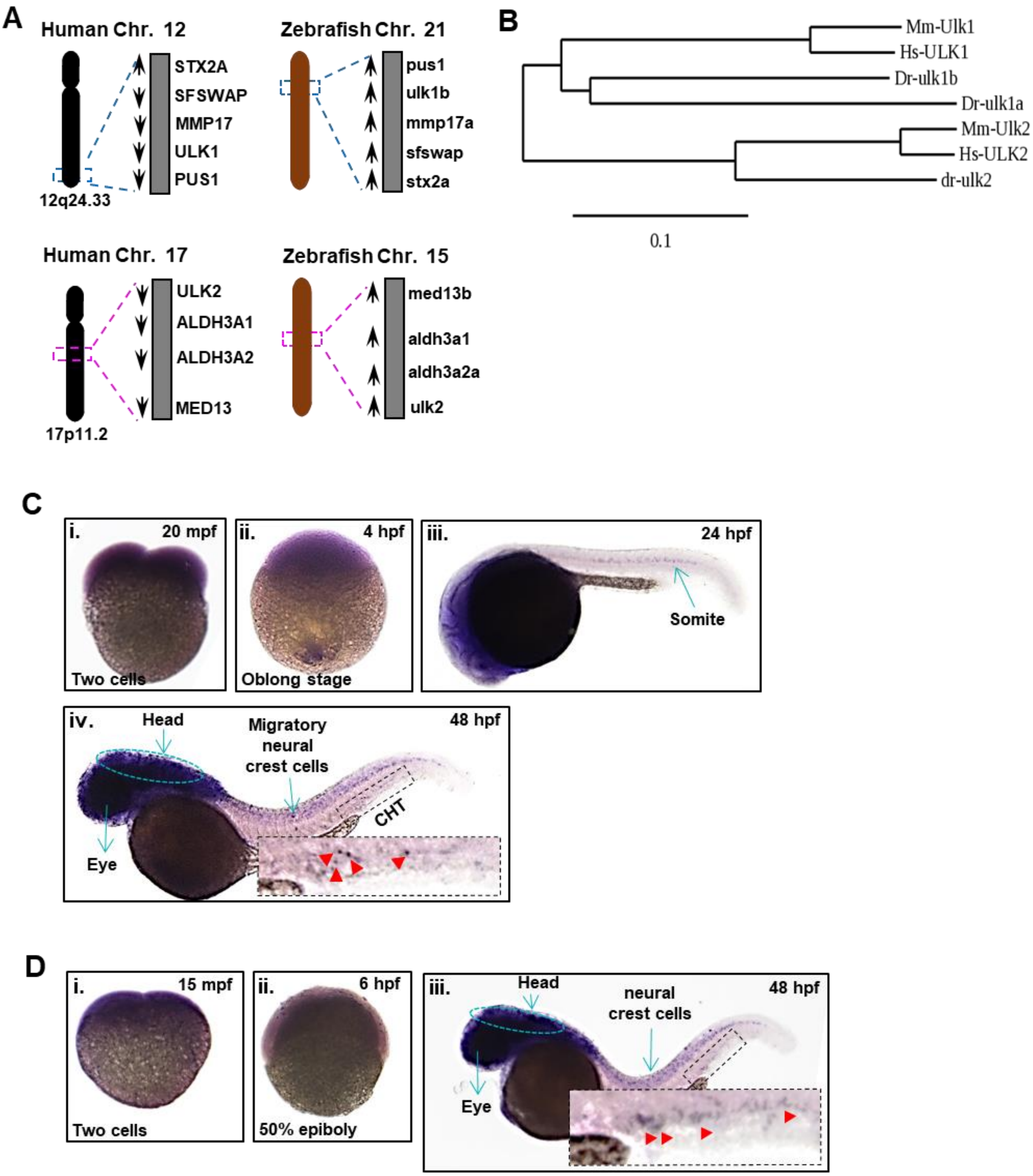

A
Human Chr. 12
STX2A
SFSWAP
MMP17
ULK1
PUS1
12q24.33
Zebrafish Chr. 21
pus1
ulk1b
mmp17a
sfswap
stx2a
Human Chr. 17
ULK2
ALDH3A1
ALDH3A2
MED13
17p11.2
Zebrafish Chr. 15
med13b
aldh3a1
aldh3a2a
ulk2
B
Mm-Ulk1
Hs-ULK1
Dr-ulk1b
Dr-ulk1a
Mm-Ulk2
Hs-ULK2
dr-ulk2
0.1
C
i. 20 mpf
Two cells
ii. 4 hpf
Oblong stage
iii. 24 hpf
Somite
iv. 48 hpf
Head
Migratory neural crest cells
CHT
Eye
D
i. 15 mpf
Two cells
ii. 6 hpf
50% epiboly
iii. 48 hpf
Head
neural crest cells
Eye

**Figure S2.**

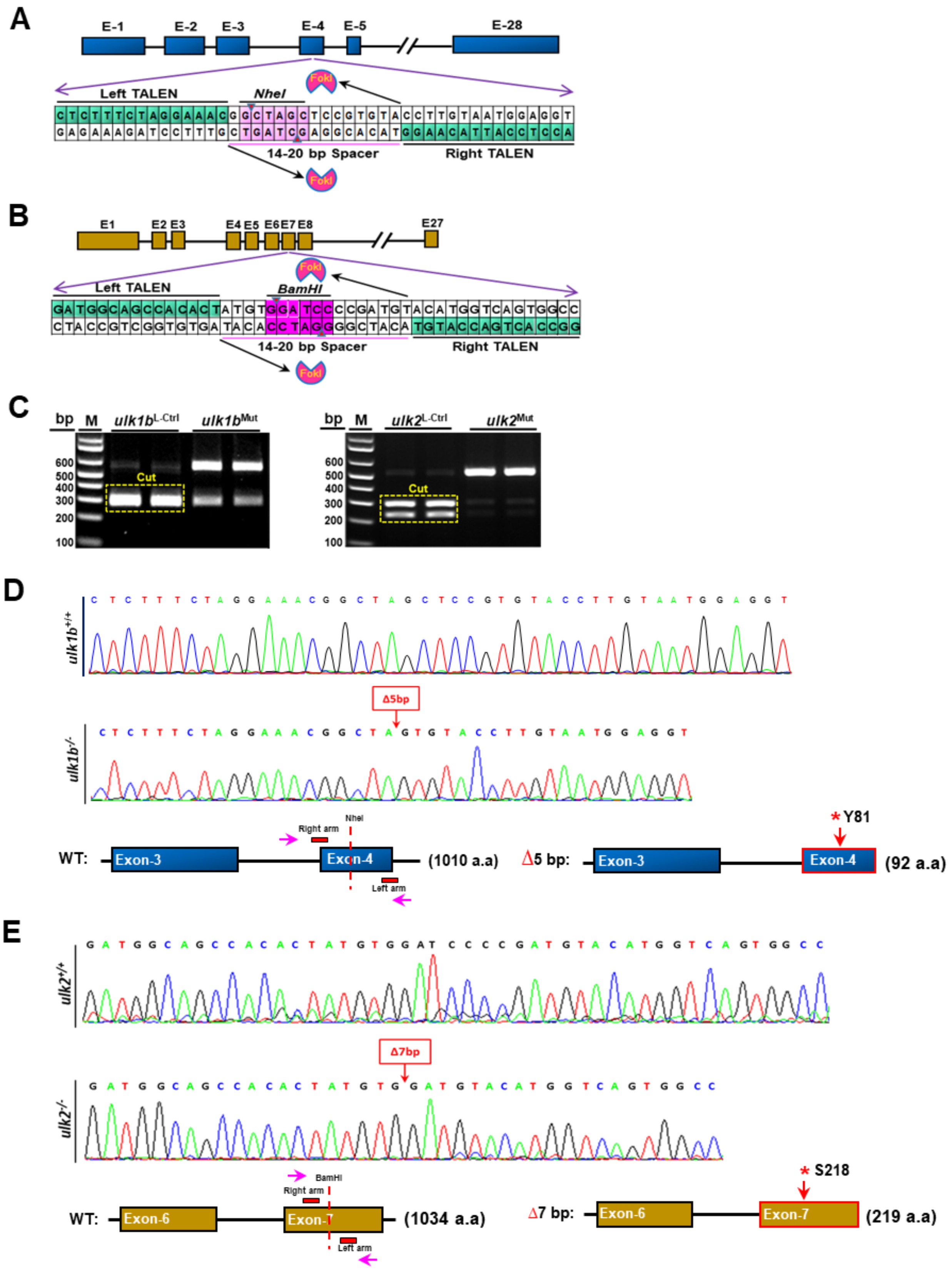

A
E-1
E-2
E-3
E-4
E-5
E-28
FokI
Left TALEN
NheI
14-20 bp Spacer
Right TALEN
B
E1
E2 E3
E4 E5 E6 E7 E8
E27
BamHI
C
bp
M
ulk1b L-Ctrl
ulk1b Mut
ulk2 L-Ctrl
ulk2 Mut
Cut
D
ulk1b+/+
ulk1b-/-
Δ5bp
WT:
Exon-3
Exon-4
(1010 a.a)
Right arm
Left arm
Δ5 bp:
*Y81
(92 a.a)
E
ulk2+/+
ulk2-/-
Δ7bp
Exon-6
Exon-7
(1034 a.a)
Δ7 bp:
*S218
(219 a.a)

**Figure S3.**

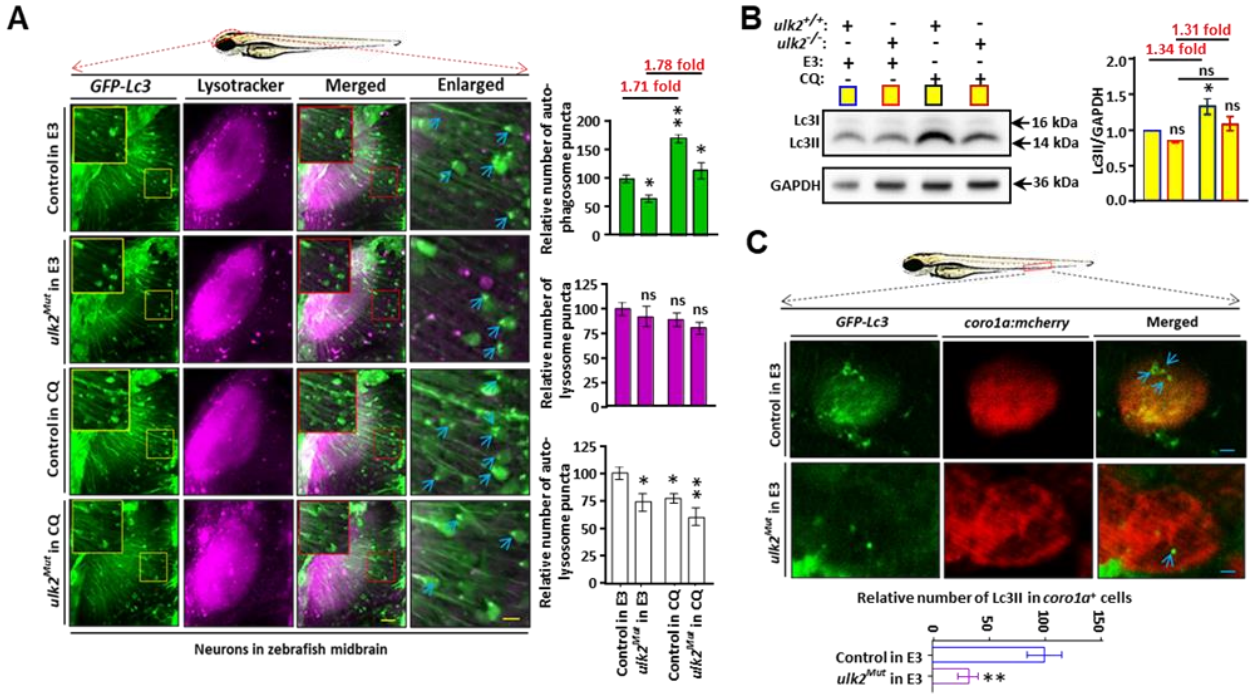
A
GFP-Lc3
Lysotracker
Merged
Enlarged
Control in E3
ulk2Mut in E3
Control in CQ
ulk2Mut in CQ
Neurons in zebrafish midbrain
1.78 fold
1.71 fold
Relative number of auto-phagosome puncta
Relative number of lysosome puncta
Relative number of auto-lysosome puncta
B
ulk2+/+:
ulk2-/-:
E3:
CQ:
Lc3I
Lc3II
GAPDH
16 kDa
14 kDa
36 kDa
1.31 fold
1.34 fold
Lc3II/GAPDH
C
GFP-Lc3
coro1a:mcherry
Merged
Relative number of Lc3II in coro1a+ cells

**Figure S4.**

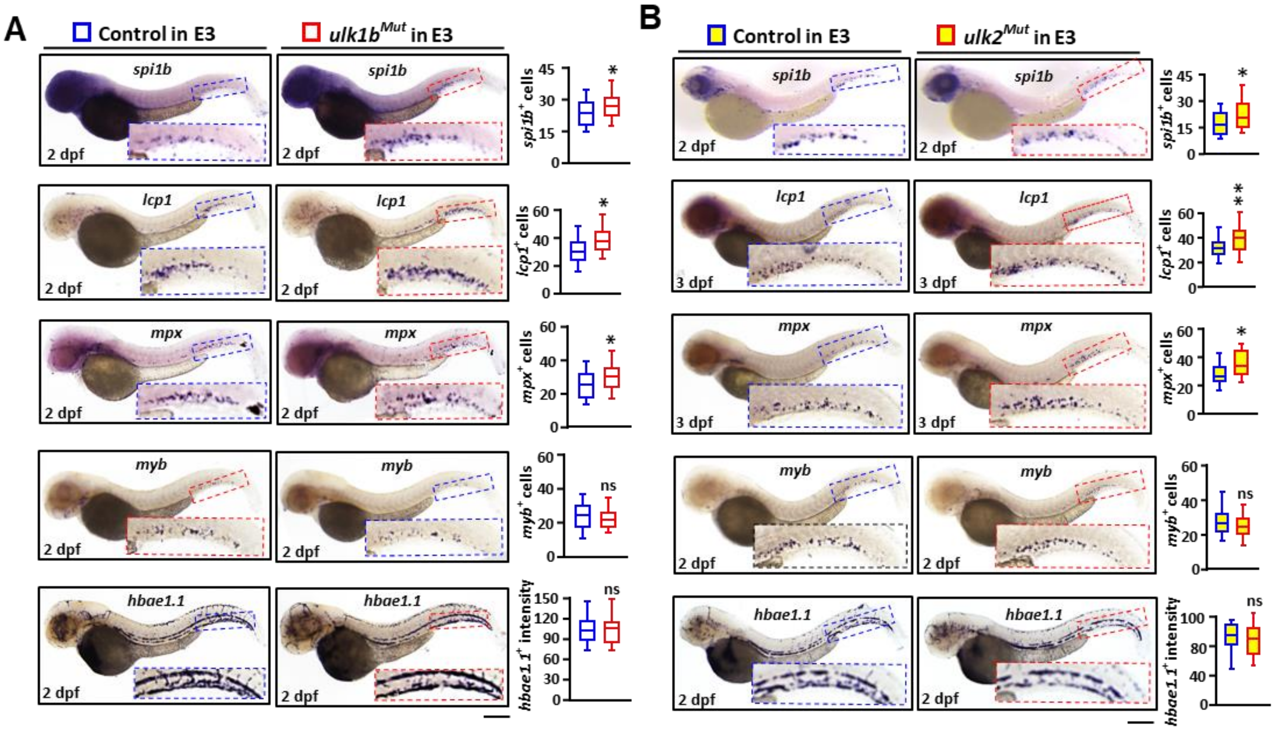
A
Control in E3
ulk1bMut in E3
spi1b
lcp1
mpx
myb
hbae1.1
2 dpf
spi1b+ cells
lcp1+ cells
mpx+ cells
myb+ cells
hbae1.1+ intensity
ns
B
Control in E3
ulk2Mut in E3
3 dpf

**Figure S5.**

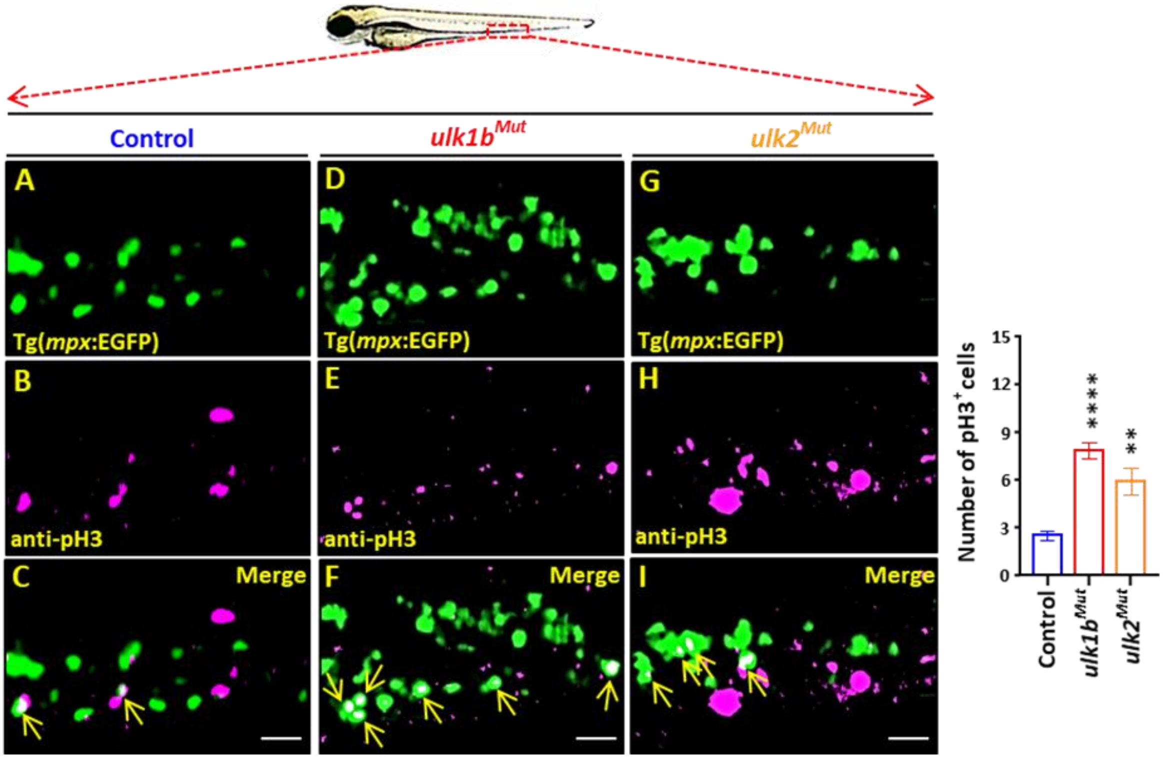
Control
ulk1b^Mut
ulk2^Mut
A
Tg(mpx:EGFP)
B
anti-pH3
C
Merge
D
Tg(mpx:EGFP)
E
anti-pH3
F
Merge
G
Tg(mpx:EGFP)
H
anti-pH3
I
Merge
Number of pH3+ cells
15
12
9
6
3
0
****
**
Control
ulk1b^Mut
ulk2^Mut

Figure S6.

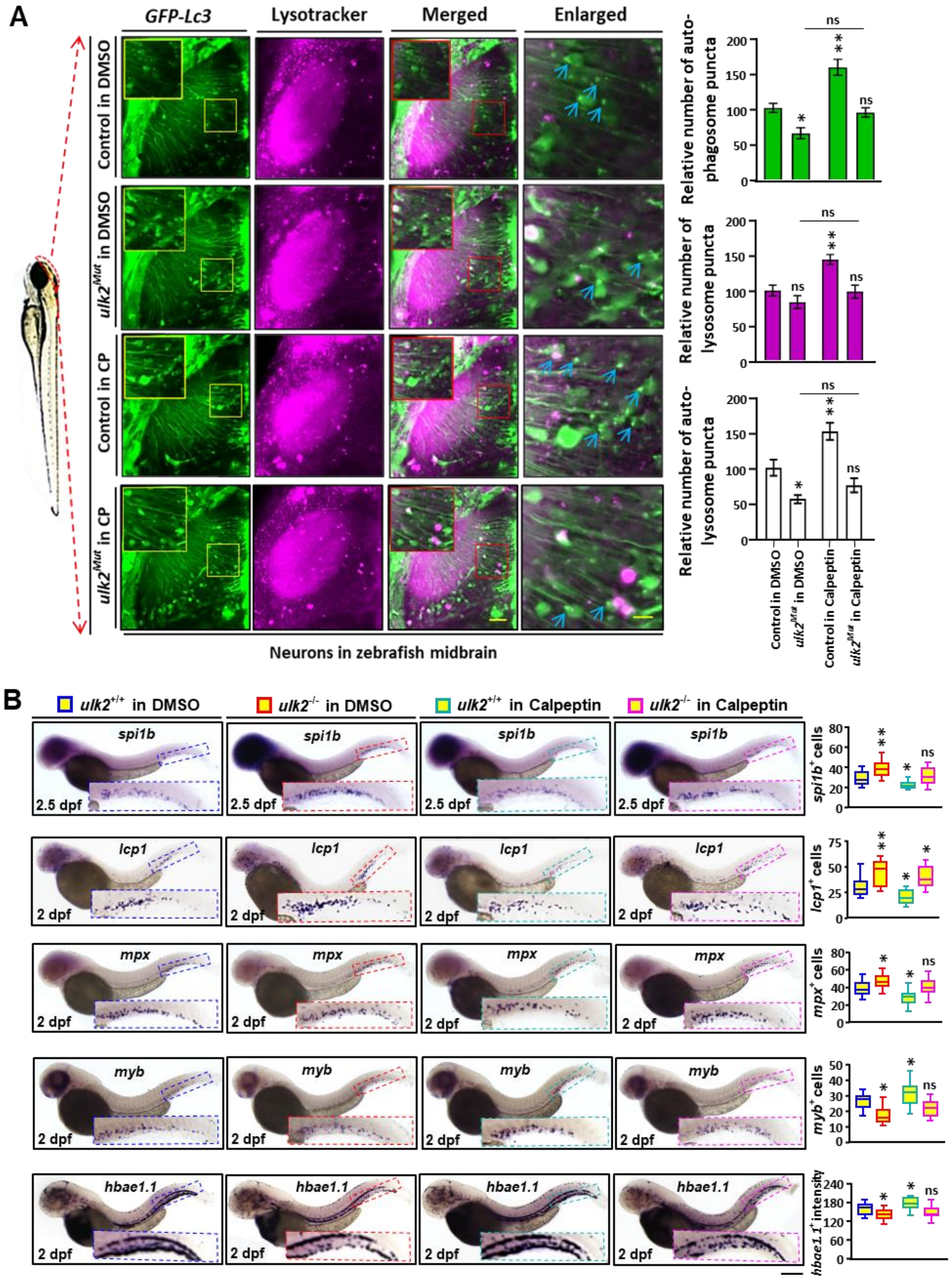
A
GFP-Lc3
Lysotracker
Merged
Enlarged
Control in DMSO
ulk2Mut in DMSO
Control in CP
ulk2Mut in CP
Neurons in zebrafish midbrain
Relative number of auto-phagosome puncta
Relative number of lysosome puncta
Relative number of auto-lysosome puncta
ns
Control in DMSO
ulk2Mut in DMSO
Control in Calpeptin
ulk2Mut in Calpeptin
B
ulk2+/+ in DMSO
ulk2-/- in DMSO
ulk2+/+ in Calpeptin
ulk2-/- in Calpeptin
spi1b
lcp1
mpx
myb
hbae1.1
2.5 dpf
2 dpf
spi1b+ cells
lcp1+ cells
mpx+ cells
myb+ cells
hbae1.1+ intensity
